# Photosystem II does not convert nascent oxygen to the poisonous singlet form

**DOI:** 10.1101/777029

**Authors:** Heta Mattila, Esa Tyystjärvi

## Abstract

In the light, the Mn_4_CaO_5_ complex of Photosystem II (PSII) splits water producing O_2_ and the triplet state of the primary donor (^3^P_680_) of PSII generates reactive singlet oxygen (^1^O_2_). We show that nascent O_2_ is not converted to ^1^O_2_, but originates exclusively from ambient O_2_, indicating that the sensitivity of PSII to oxidative damage is not a consequence of the water-splitting *per se*, and showing that the suggested oxygen channels function nearly perfectly, conveying nascent O_2_ out of the reach of ^3^P_680_. This may have been crucial during evolution of oxygenic photosynthesis, as ^3^P_680_ cannot be quenched by carotenoids that protect non-oxygenic photosystems. In addition, the data indicate that a ^1^O_2_-independent mechanism contributes to the light-induced damage of PSII.

## Introduction

Cyanobacteria, algae and plants harvest light energy from the Sun, supporting life on the Earth. Photosystem II (PSII), the unique water splitting protein machine of photosynthesis, converts light energy to a chemical form and starts the photosynthetic electron transfer chain. Excitation of one of the PSII reaction center chlorophylls (Chls), called P_680_, rapidly leads to reduction of a primary electron acceptor, a pheophytin molecule^1^. The strongest known biological oxidant, P680^+^, formed by the charge separation, then extracts an electron from the oxygen evolving Mn_4_CaO_5_ complex (OEC) that catalyzes oxidation of water and the associated formation of molecular oxygen.

Use of light energy for water splitting comes with a challenge. An excited chlorophyll molecule may assume a triplet spin configuration (^3^Chl) which readily donates energy to ground state O_2_ which itself is a triplet. The reaction produces a ground state Chl and a singlet excited state of O_2_^2^. Singlet O_2_ can occur in two forms but 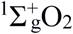, which has the two outermost electrons on two different orbitals, rapidly decays to ^1^Δ_g_O_2_ (abbreviated as ^1^O_2_ here) that has both electrons on the same orbital. ^1^O_2_ is highly reactive, damages lipids and proteins^3^, and is considered to be the most dangerous reactive oxygen species (ROS) occurring in plants^4. 1^O_2_ produced by an added sensitizer chemical is also known to damage PSII^5^. The ^3^Chl required for ^1^O_2_ formation may be formed either in the reaction center or by intersystem crossing among the light harvesting antenna complexes^6–7^. In the reaction center, triplets are mainly formed by charge recombination reactions, which are back reactions of photosynthetic electron transfer^8–9^.

Chls of the light harvesting complexes are efficiently protected against oxidation by ^1^O_2_ by carotenoids^10–11^ that quench both ^3^Chl and ^1^O_2_. Carotenoids also protect the reaction centers of non-oxygenic photosynthetic bacteria in which bacteriochlorophyll would otherwise produce ^1^O_2_^12^. In addition, carotenoids may protect the reaction center of photosystem I (PSI)^13^. P_680_^+^ is, however, so oxidizing that it would irreversibly oxidize a carotenoid located close enough to physically quench ^3^P_680_^14^. Therefore, formation of ^3^P_680_ leads to formation of ^1^O_2_ whenever a ground-state O_2_ molecule is close by. Accordingly, production of ^1^O_2_ has been measured from cyanobacteria and plants in the light, both *in vitro* and *in vivo*^14–17^. Correlation of the amount of ^1^O_2_ with decay kinetics of ^3^P_680_^18^ and accumulation of β- carotene endoperoxide during high light stress^19^ strongly suggest that most of this ^1^O_2_ is produced by ^3^P_680_.

The unavoidable formation of ^3^P_680_ and the resulting high probability of formation of ^1^O_2_ from O_2_ poses a dilemma, as OEC that produces an O_2_ molecule per every fourth photon absorbed by PSII^1^, resides only ∼20 Å from P_680_ ^20^, potentially exposing P_680_ to a high oxygen pressure. Can O_2_ produced by OEC instantly diffuse to P_680_, become converted to ^1^O_2_ and possibly damage PSII? To answer this question, we measured the production of ^1^O_2_ by isolated thylakoid membranes in strong light using a histidine-based method^18,21^. Membrane inlet mass spectroscopy (MIMS) was used for the analysis of gas exchanges, allowing us to independently monitor the fates of ambient O_2_ and the nascent O_2_ produced by water splitting in PSII.

## Materials and Methods

### Plant material

Pumpkin (*Cucurbita maxima* L.) was grown at the photosynthetic photon flux density (PPFD) of 200 μmol m^-2^s^-1^, with a 16 h/8 h light/dark period at 20 °C. For isolation of thylakoid membranes, leaves were ground in ice-cold buffer (40 mM HEPES (pH 7.4), 0.3 M sorbitol, 10 mM MgCl_2_, 1 mM ethylenediamine tetra-acetic acid, 1 M glycine betaine and 1% bovine serum albumin), filtered and centrifuged (5 min; 1100 x g). The pellet was resuspended in osmotic shock buffer (10 mM HEPES (pH 7.4), 5 mM sorbitol and 10 mM MgCl_2_), centrifuged (5 min; 2000 x g) and stored at −75 °C in a storage buffer (10 mM HEPES (pH 7.4), 0.5 M sorbitol, 10 mM MgCl2 and 5 mM NaCl). The Chl content of the thylakoids was quantified spectrophotometrically^22^.

### O_2_ measurements

O_2_ was measured either with an O_2_ electrode (Hansatech, King’s Lynn, UK) or with MIMS. MIMS was operated as follows: O_2_ isotopes (^16^O_2_ and ^18^O_2_) were measured with Prima PRO Process Mass Spectrometer (Thermo Scientific(tm)) connected with vacuum lines to the sample chamber (modified from Hansatech Instruments Ltd O_2_ electrode chamber). The chamber was separated with a PTFE membrane (Hansatech Instruments Ltd, UK). Air was removed from the sample (1 ml) with nitrogen flow and replaced with ^18^O_2_ (adjusted to about 100 µM; approximately 30 µM ^16^O_2_ remained). Measurements were started after the gases in the sample reached an equilibrium. Two-point calibration was done by flushing the sample first with nitrogen (0 µM O_2_) and then with air (O_2_-saturated, i.e. 253 µM O_2_; ^18^O_2_ was assumed to have similar response as ^16^O_2_). Rates of O_2_ consumption or evolution were calculated at 25–72 s after switching on the light, and slow thylakoid-independent decrease in O_2_ concentration in the chamber during the measurement, due to diffusion to the vacuum lines, was assumed to be linear during the 150 s measurement and was subtracted from the calculated values.

### ^1^O_2_ measurement

^1^O_2_ produced in high light (PPFD 3000 µmol m^-2^s^-1^ of white light) by thylakoid membranes (100 µg chl/ml) was measured in photoinhibition buffer (40 mM HEPES-KOH (pH 7.4), 1 M betaine monohydrate, 330 mM sorbitol, 5 mM MgCl_2_ and 5 mM NaCl) at 22 or 25 °C, as indicated, by following the consumption of O_2_ by 20 mM histidine^18,21^.

### High-light treatments

Isolated pumpkin thylakoids in the photoinhibition buffer, or lincomycin-treated leaves, were illuminated 0–180 min at 20 °C with white light (PPFD 2000 µmol m^-2^s^-1^). Prior to the illumination, excised pumpkin leaves were incubated over-night in low light with the petioles in lincomycin (0.4 mg/ml) solution. The activity of PSII was measured by illuminating thylakoid membranes with saturating light in the presence of 0.5 mM 2,6-dimethylbenzoquinone at 22 °C in buffer (40 mM HEPES-KOH (pH 7.6), 1 M betaine monohydrate, 330 mM sorbitol, 5 mM MgCl_2_, 5 mM NaCl, 1 mM KH_2_PO_4_ and 5 mM NH_4_Cl).

## Results & Discussion

### Nascent oxygen is not involved in the production of singlet oxygen

Thylakoid membranes, isolated from pumpkin leaves, were illuminated for 150 s in strong light in the presence or absence of 20 mM histidine, and changes in the concentrations of two oxygen isotopes (^16^O_2_ and ^18^O_2_) were measured with MIMS. Histidine is an efficient chemical scavenger of ^1^O_2_ but does not react rapidly with ground-state O_2_ or other ROS^21^, and therefore the loss of O_2_ in the presence of histidine measures the formation of ^1^O_2_. Before the measurements with MIMS, ∼80 % of the dissolved O_2_ in the sample buffer was replaced with the heavier isotope (^18^O_2_). In this way, ambient ^18^O_2_ was distinguished from ^16^O_2_ that originated from oxidation of water.

When strong light was switched on (PPFD 3000 µmol m^-2^s^-1^), evolution of ^16^O_2_, corresponding to water splitting by OEC, was observed (Fig. 1A), but the concentration of the ambient ^18^O_2_ decreased at a higher rate (Fig. 1B), which led to net oxygen consumption by the rate of 2.3 µmol O_2_ mg Chl^-1^ h^-1^ (Fig. 1C). The result is expected, as isolated thylakoid membranes do not fix CO_2_ but instead reduce O_2_, mostly at PSI^23^, with possible smaller contributions by plastoquinone pool, plastid terminal oxidase or PSII.

**Figure 1.**
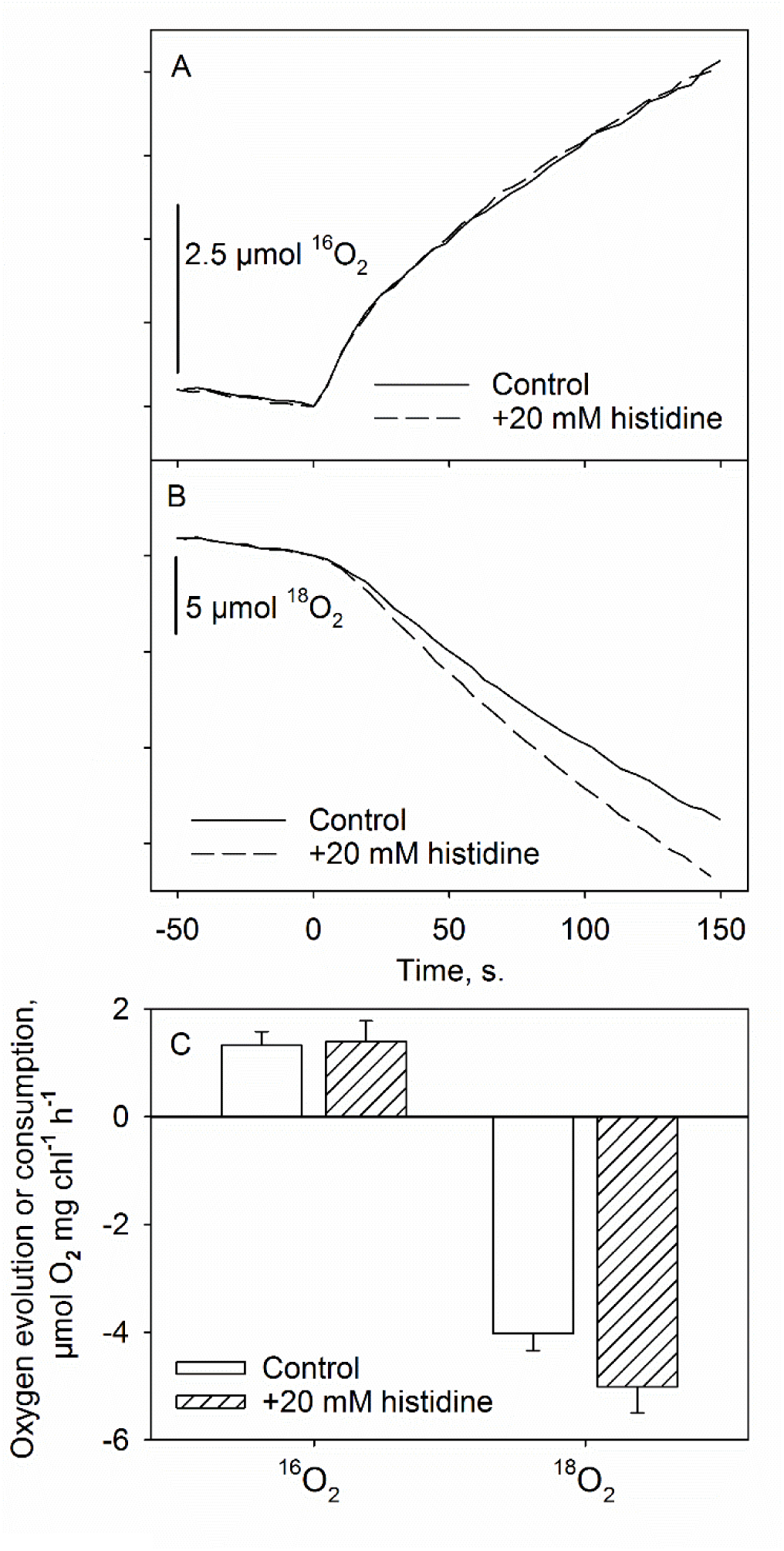
Changes in concentrations of ^16^O_2_ and ^18^O_2_ during incubation of pumpkin thylakoids at 25 °C in the absence or presence of histidine, measured with MIMS. The curves have been normalized to the values at the time point 0 at which strong light was switched on. In the beginning, the control sample contained 30.8 (±5.1) µM ^16^O_2_ and 107.5 (±19.6) µM ^18^O_2_, and the histidine sample contained 31.2 (±9.5) µM ^16^O_2_ and 117.1 (±49.6) µM ^18^O_2_. The rates of O_2_ evolution or consumption (C) have been calculated from (A) and (B). All data represent mean values from six independent experiments, and standard deviations (s.d.) are shown as error bars in (C). The difference between the rates of ^16^O_2_ evolution was not statistically significant whereas the increase in the rate of consumption of ^18^O_2_ was significant (P = 0.0027; students t-test).

More interestingly, the addition of histidine did not affect the rate of ^16^O_2_ evolution (Figs. 1A and 1C) while the consumption of ambient ^18^O_2_ was significantly accelerated (Figs. 1B–C). The result shows that the ^1^O_2_ that reacted with the added histidine derived exclusively from the ambient ^18^O_2_ whereas nascent ^16^O_2_, originating from water splitting, did not contribute to ^1^O_2_ production. The presence of ∼20 % of ^16^O_2_ in the ambient air leads to a slight underestimation of the O_2_ evolution rate, as some ^16^O_2_ was simultaneously consumed by reduction of oxygen at PSI. This, however, does not affect the conclusion about the fate of the nascent O_2_.

### Water splitting activity of PSII is not required for singlet oxygen production

The result raises the question whether capacity to evolve O_2_ is required for ^1^O_2_ production at all. For this, we compared the rate of ^1^O_2_ production with the oxygen evolving activity of PSII in thylakoid membranes. To obtain membranes with different activities, thylakoids were illuminated for 0–180 min with strong light. In addition, pumpkin leaves were subjected to similar illumination treatments and thylakoid membranes were isolated for the ^1^O_2_ production assay from the treated leaves; lincomycin pre-treatment prevented recovery of PSII in the illuminated leaves.

The data show that the rate of ^1^O_2_ production did not depend on the O_2_ evolving activity of PSII (Fig. 2). Thus, nascent O_2_ is not converted to ^1^O_2_, and the ability to produce O_2_ is not needed for ^1^O_2_ production. The result is in line with the findings that the rate of ^1^O_2_ production remains unchanged for 300 min under high-light illumination of *Arabidopsis* leaves^17^ and that the D1 protein of PSII reaction center may not be required for ^1^O_2_ production^24^. However, Hideg et al.^16^ calculated that the amount of inactive PSII centers with the D1 protein still present correlated with the rate of ^1^O_2_ production. The reasons for the contradictory results may stem from the fact that we recorded instantaneous ^1^O_2_ production whereas Hideg et al.^16^ used a cumulative method.

**Figure 2.**
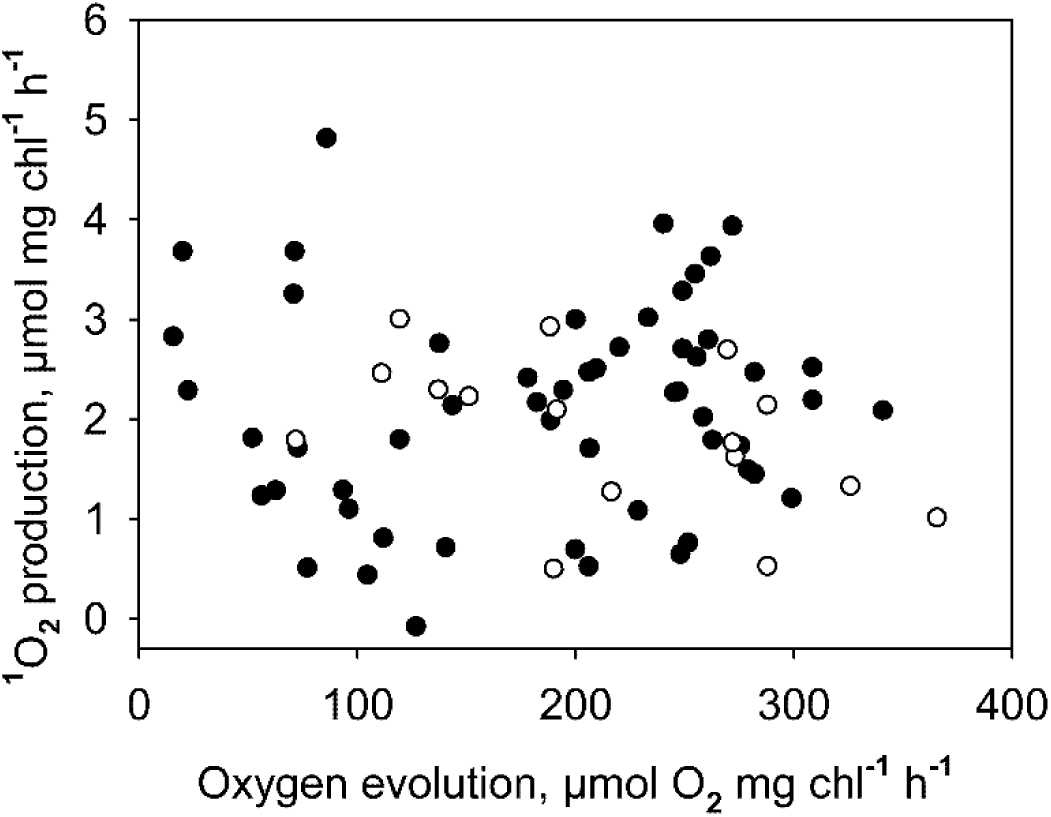
^1^O_2_ production in photoinhibited thylakoids. Relationship between the light-saturated O_2_ evolution activity of PSII (measured in the presence of an artificial electron acceptor) and production of ^1^O_2_ (separately measured as histidine-mediated O_2_ consumption) was measured with an oxygen electrode at 22 °C from isolated pumpkin thylakoid membranes. Prior to the measurement, thylakoids (solid dots) or lincomycin-treated pumpkin leaves (open dots) were illuminated for 0–180 min at PPFD 2000 µmol m^-2^s^-1^. When leaves were used, thylakoid membranes were isolated directly after the illumination treatment and subjected to the ^1^O_2_ production and O_2_ evolution assays.

### The reaction center of PSII is protected from nascent oxygen

Our results show that O_2_ produced by the OEC cannot be instantly converted to ^1^O_2_. Of course, practically all free O_2_ originates from water, oxidized by PSII, and eventually (time > 150 s, as used in the present study) nascent O_2_ will became ambient O_2_. The present results indicate, however, that the O_2_ production *per se* do not render PSII more vulnerable to oxidative damage than any other photosynthetic protein complex.

How is the reaction center of PSII protected from the nascent O_2_? Both experimental data^25^ and structural analysis^26^ have suggested that PSII has protein channels that divert nascent O_2_ out of the OEC. O_2_ may exit from PSII into the thylakoid lumen, possibly close to the membrane surface^26^ or into the thylakoid membrane^25^. Our results indicate that at least one of the proposed channels functions efficiently, as nascent O_2_ does not have a direct access to the reaction center of PSII.

Minimizing ^1^O_2_ production was extremely important during the early evolution of oxygenic bacteria, when the antioxidative protective mechanisms were not yet fully functional^27^. Despite the O_2_ evolution by PSII, the O_2_ concentration of a cyanobacterial cell rapidly equilibrates with the environment, and consequently the O_2_ concentration inside a bacterial cell is only little higher than the ambient O_2_ concentration^28^. Thus, prevention of direct contact of nascent O_2_ with P_680_ led to full avoidance of ^1^O_2_ formation in the anoxic or micro-aerobic environment of early cyanobacteria. It has been speculated that avoiding the contact of O_2_ with the reaction center would be important to protect PSII even in extant oxygenic organisms^29^.

### A singlet-oxygen-independent mechanism must be involved in photoinhibition

PSII is continuously damaged in the light by a reaction known as photoinhibition^30–32^, and evidence has been presented for both a mechanism based on oxidation by ^1^O_2_^21^ and for direct light-induced damage to the OEC^33^. The finding that nascent O_2_ is not converted to ^1^O_2_ indicates that a ^1^O_2_-dependent photoinhibition mechanism would be strongly dependent on ambient O_2_. Earlier data, however, show that ambient O_2_ alleviates photoinhibition in oxygen evolving PSII particles^34^ or exacerbates photoinhibition in spinach leaves by a small amount^35^. Thus, a ^1^O_2_ independent mechanism must strongly contribute to photoinhibition of PSII in visible light.

## Acknowledgments

Dr. Duncan Fitzpatrick is thanked for generous technical and scientific advice, prof. Maarit Karonen for fruitful discussions and Sofia Vesterkvist for assistance in experiments. Turku University Foundation (12353), Vilho, Yrjö and Kalle Väisälä Foundation and Academy of Finland (307335) are thanked for financial support.

## Author contributions

ET conceived and supervised the study; HM designed and performed experiments; HM wrote the manuscript with contribution from ET.

## Data Availability

Raw data can be found in https://seafile.utu.fi/d/66f0a45946b843228f41/.

## Abbreviations

^1^O_2_: singlet oxygen (^1^Δ_g_O_2_);
^3^Chl: triplet chlorophyll;
Chl: Chlorophyll;
MIMS: membrane inlet mass spectroscopy;
OEC: oxygen evolving complex;
PPFD: photosynthetic (400–700 nm) photon flux density;
PSI: Photosystem I;
PSII: Photosystem II;
P_680_: reaction center chlorophylls (the primary donor) of PSII;
ROS: reactive oxygen species

